# Genomic regression of claw keratin, taste receptor and light-associated genes inform biology and evolutionary origins of snakes

**DOI:** 10.1101/127654

**Authors:** Christopher A. Emerling

## Abstract

Regressive evolution of anatomical traits corresponds with the regression of genomic loci underlying such characters. As such, studying patterns of gene loss can be instrumental in addressing questions of gene function, resolving conflicting results from anatomical studies, and understanding the evolutionary history of clades. The origin of snakes coincided with the regression of a number of anatomical traits, including limbs, taste buds and the visual system. By studying the genomes of snakes, I was able to test three hypotheses associated with the regression of these features. The first concerns two keratins that are putatively specific to claws. Both genes that encode these keratins were pseudogenized/deleted in snake genomes, providing additional evidence of claw- specificity. The second hypothesis is whether snakes lack taste buds, an issue complicated by unequivocal, conflicting results in the literature. I found evidence that different snakes have lost one or more taste receptors, but all snakes examined retained at least some capacity for taste. The final hypothesis I addressed is that the earliest snakes were adapted to a dim light niche. I found evidence of deleted and pseudogenized genes with light-associated functions in snakes, demonstrating a pattern of gene loss similar to other historically nocturnal clades. Together these data also provide some bearing on the ecological origins of snakes, including molecular dating estimates that suggest dim light adaptation preceded the loss of limbs.

## 1. Introduction

The evolutionary origin and diversification of lineages is frequently attributed to key innovations of traits, but regressive evolution (Fong et al., 1995), the loss of formerly adaptive traits through evolutionary time, can also be a powerful force in the shaping of clades (Albalat and Cañestro, 2016). As phenotypic traits regress in a species, their genomes are expected to retain a corresponding signature, including the retention of unitary pseudogenes and whole gene deletions (Meredith et al., 2009; Emerling and Springer, 2014; Feng et al., 2014; Albalat and Cañestro, 2016). This allows researchers to test hypotheses of gene function, clarify phenotypes contradicted by different reports, and uncover aspects of a lineage’s evolutionary history. Additionally, the retention of loss-of-function mutations shared by multiple species can help index the timing of genomic and phenotypic regression, while modeling DNA sequence evolution in pseudogenes can provide additional resolution, thereby helping to understand the sequence of events in a clade’s evolutionary history.

Here I test hypotheses of gene functionality, phenotype and evolutionary history associated with the origin of snakes (Serpentes). Adaptations in the earliest snakes involved the regression of a number of anatomical traits present in the ancestral squamate, including limbs, taste buds and portions of the eye involved in bright light (photopic) vision. The loss of limbs co-occurred with the loss of claws, and potentially their associated keratins associated. Previous research has implicated two keratin genes (*HA1*, *HA2*) as claw-specific (Eckhart et al., 2008; Alibardi et al., 2011; Dalla Valle et al., 2011), but the data are quite limited in taxonomic breadth. Given that snakes lack claws, finding pseudogenic and/or deleted copies of these genes would provide strong support for the claw-specificity hypothesis. If correct, these loci could then be used as a genomic proxy to test for patterns and timing of leg loss across squamates. The loss of taste in snakes is controversial, with some studies finding evidence for taste buds (Kroll, 1973; Berkhoudt et al., 2001), and the most extensive comparative study unequivocally concluding that taste buds are completely absent (Young, 1997). Given that the genomic basis for taste is well understood (Chandrashekar et al., 2006; Yarmolinsky et al., 2009), the presence or absence of associated taste genes should be able to weigh in on these competing hypotheses. Finally, serpent eye anatomy is strongly suggestive of an early period of dim light (scotopic) adaptation in snake history (Walls, 1942). Various genes encoding visual and non-visual photopigments, as well as enzymes involved in reducing ultraviolet (UV) photo-oxidative damage, have a clear functional association with light (Bellingham et al., 2006; Frigato et al., 2006; Su et al., 2006; Tomonari et al., 2008; Davies et al., 2012; Ohuchi et al., 2012; Wada et al., 2012; Osborn et al., 2015). Species occupying dim light for extensive periods of time are predicted to undergo regression of such loci, since maintaining the expression of such proteins would be energetically costly if insufficient light is available to induce a physiological response. Finding pseudogenes and/or gene deletions of such light-associated genes in all snakes, especially with shared inactivating mutations, would corroborate the hypothesis that the earliest snakes were indeed dim light adapted.

These regressed traits can also weigh in on a vigorous debate concerning the origins of snakes. The ecological context for serpent origins has been attributed to early marine (Caldwell and Lee, 1997; Lee et al., 1999; Caprette et al., 2004; Lee, 2005; Lee et al., 2016), fossorial (Walls, 1942; Apesteguía and Zaber, 2006; Longrich et al., 2012; Yi and Norell, 2015; Martill et al., 2015), and/or nocturnal adaptations (Hsiang et al., 2015; reviewed in Simões et al., 2015). These models are variably consistent with the loss of limbs, taste buds, and photopic photoreception, and the timing and distribution of gene loss may have some bearing on these hypotheses.

## 2. Methods

### 2.1. Taxonomic sampling

Serpentes is generally divided into two infraorders: Scolecophidia (blind snakes) and Alethinophidia (all remaining snakes). Here I focused on the publicly-available genomes of eight species of alethinophidian snakes: a pythonid (*Python bivittatus* [Burmese python]) and seven colubroids (*Crotalus mitchellii* [speckled rattlesnake; Viperidae], *Crotalus horridus* [timber rattlesnake; Viperidae], *Protobothrops mucrosquamatus* [brown spotted pit viper; Viperidae], *Vípera berus* [European adder; Viperidae], *Ophiophagus hannah* [king cobra; Elapidae], *Thamnophis sirtalis* [common garter snake; Colubridae], *Pantherophis guttatus* [corn snake; Colubridae]; electronic supplementary material [ESM], Table S1; Castoe et al., 2011; Vonk et al., 2013; Gilbert et al., 2014; Ullate-Agote et al., 2014; Castoe et al., 2014). Though this prevented confirmation of gene losses common to both infraorders of snakes, gene inactivation dating estimates (see below) predict that some gene losses occurred in the stem serpent lineage. I made additional comparisons to outgroup taxa to determine whether the gene losses were unique to snakes, with special focus on the nocturnal *Gekko japonicus* (Schlegel’s Japanese gecko) for light-associated genes.

### 2.2. Determining gen e functionality

To collect data on the functionality of the 27+ claw keratin, gustatory and light-associated genes examined in this study, I used a combination of NCBI’s Eukaryotic Genome Annotation (EGA) pipeline and direct queries of NCBI’s Whole Genome Shotgun contig database (WGS). To utilize EGA, I BLASTed (discontiguous megablast) reference sequences against NCBI’s nucleotide collection using the taxon ID “Squamata” and recorded annotations of “PREDICTED” sequences. At the time this study was performed, EGA was only available for five squamates: *Python bivittatus*, *Protobothrops mucrosquamatus*, *Thamnophis sirtalis*, *Gekko japonicus* and *Anolis carolinensis*. EGA provides evidence of gene non-functionality in two ways: negative BLAST results, due to absence of the gene in the assembly or extensive sequence divergence, as is typical of many pseudogenes, and annotation of a predicted transcript as a “Low quality protein”. The latter is indicated if the predictive model replaces nucleotides in premature stop codons and/or frameshift indels in the original genome assembly. If neither of these occurred, I assumed that the gene is functional.

When I found evidence of gene dysfunction, I directly queried NCBI’s Whole Genome Shotgun Contig database (WGS) using *in silico* gene probes. These probes were designed by BLASTing (megablast) a reference coding sequence (e.g., mRNA) against a reference genome (typically *Anolis carolinensis’*) and aligning the coding sequence to the relevant contig(s) using MUSCLE (Edgar, 2004) in Geneious ver. 9.1.2 (Kearse et al., 2012). The resulting probe encompassed the coding sequence, introns and flanking sequences of the gene of interest. This probe was then BLASTed (blastn) against each of the focal species. In instances of negative BLAST results, I BLASTed individual exons + flanking sequence, since BLASTing of the entire probe frequently biased hits towards transposable elements located in the introns of the probe sequence. If no coding sequence was obtained, I used synteny information from outgroup species genomes in Ensembl to determine the genes flanking the locus of interest. I subsequently BLASTed reference sequences of these genes against WGS, which allowed me to determine if the flanking sequences are retained on the same contig, thereby providing evidence of whole gene deletion. After obtaining sequences from WGS, I aligned them to the probe sequence using MUSCLE in Geneious ver. 9.1.2, followed by manual adjustment. I then examined the sequences for inactivating mutations (premature stop codons, frameshift indels, and splice site mutations).

### 2.3. Gene inactivation dating analyses

In instances where inactivating mutations were discovered in one or more taxon, I performed gene inactivation dating analyses utilizing the method described in Meredith et al. (2009) with PAML ver. 4.8 (Yang, 2007). I added published sequences from GenBank and genome-derived sequences from Emerling (2016) to provide background ratio branches, and I removed ambiguous portions of the alignment before analyses. Squamate topologies were derived from Zheng and Wiens (2016). I estimated dN/dS ratios for pseudogene category branches (i.e., branches on which the gene is expected to be evolving neutrally) and then compared them to models where the pseudogene ratio was fixed at 1. If the difference was non-significant, I assumed the pseudogene ratio was 1. In only one case (*SWS2* snakes, cf1 [codon frequency model 1]) was the difference between estimated and fixed pseudogene ratios significant, so in this instance I used the estimated ratio to calculate the gene inactivation date. dN/dS ratios that exceeded 1 on mixed branches were fixed at 1 in gene inactivation date calculations, leading to the assumption that the gene was lost immediately after cladogenesis. For dN/dS ratio analyses of *TAS1R2*, I removed *Thamnophis* due to insufficient sequence overlap. Gene inactivation estimates are highly sensitive to the timetree used, and recent timetree analyses vary widely in the ages of clades relevant to this study. For example, Zheng and Wiens (2016) summarize divergence date estimates from four recent studies and show that the ages of Serpentes (113-131.1 Ma) and especially Toxicofera (140.8-184.6 Ma) are quite disparate. I used the divergence dates estimated by Zheng and Wiens (2016) for this study due to the unparalleled size of their dataset (52 genes, 4162 species, 13 fossil calibrations) and because their estimated age for the split between Serpentes and other toxicoferans (184.6 Ma) is consistent with the recent discovery of the stem serpent *Eophis* from the Middle Jurassic (Bathonian: 168.3-166.1 Ma; Caldwell et al., 2015).

### 2.4. Gene sampling

The genes included in this study are two putatively claw-specific keratin genes (*HAÏ*, *HA2*), five+ genes associated with taste (*TAS1R1*, *TAS1R2*, *TAS1R3*, *TAS2Rgene* family, *PKD2LT*), and 20 genes encoding proteins with light-dependent functions (*SWS1*, *SWS2 RH1*, *RH2*, *LWS*, *OPNP*, *OPNPT*, *OPNPP*, *OPNVA*, *OPN3*, *OPN4M*, *OPN4X*, *OPN5*, *OPN5L1*, *OPN5L2*, *RGR*, *RRH*, £V-like, *MT-Ox*, photolyase; ESM Tables S2 and S3). *TAS2Rs* vary in copy number in vertebrates, so I only utilized EGA to quantify copy number. I obtained information on *TAS2R* copy number by searching NCBI’s nucleotide collection for genes annotated as “taste receptor type 2” with *“Python"*, “*Protobothrops* and “ *Thamnophisl’* amended to respective searches.

## 3. Results

### 3.1. Claw keratins

I recovered exon 1 of the putative claw keratin gene *HA1* in *Python bivittatus*, *Ophiophagus hannah* and all four of the viperids in this study. Numerous inactivating mutations and the failure to recover exons 2-7 indicate that *HA1* is a pseudogene in all six of these species (ESM, Table S2, Dataset S1), and an 8-bp insertion shared by these snakes (ESM, Table S2) suggests that this gene was inactivated in their most recent common ancestor (MRCA). By contrast, *HA1* is intact in *Gekko japonicus*. I estimated that *HA1* was inactivated in a crown snake (Zheng and Wiens, 2016) approximately 113 Ma (point estimate; range: 108-118.5 Ma; Fig. 1; ESM, Tables S4 and S5).

**Figure 1.**
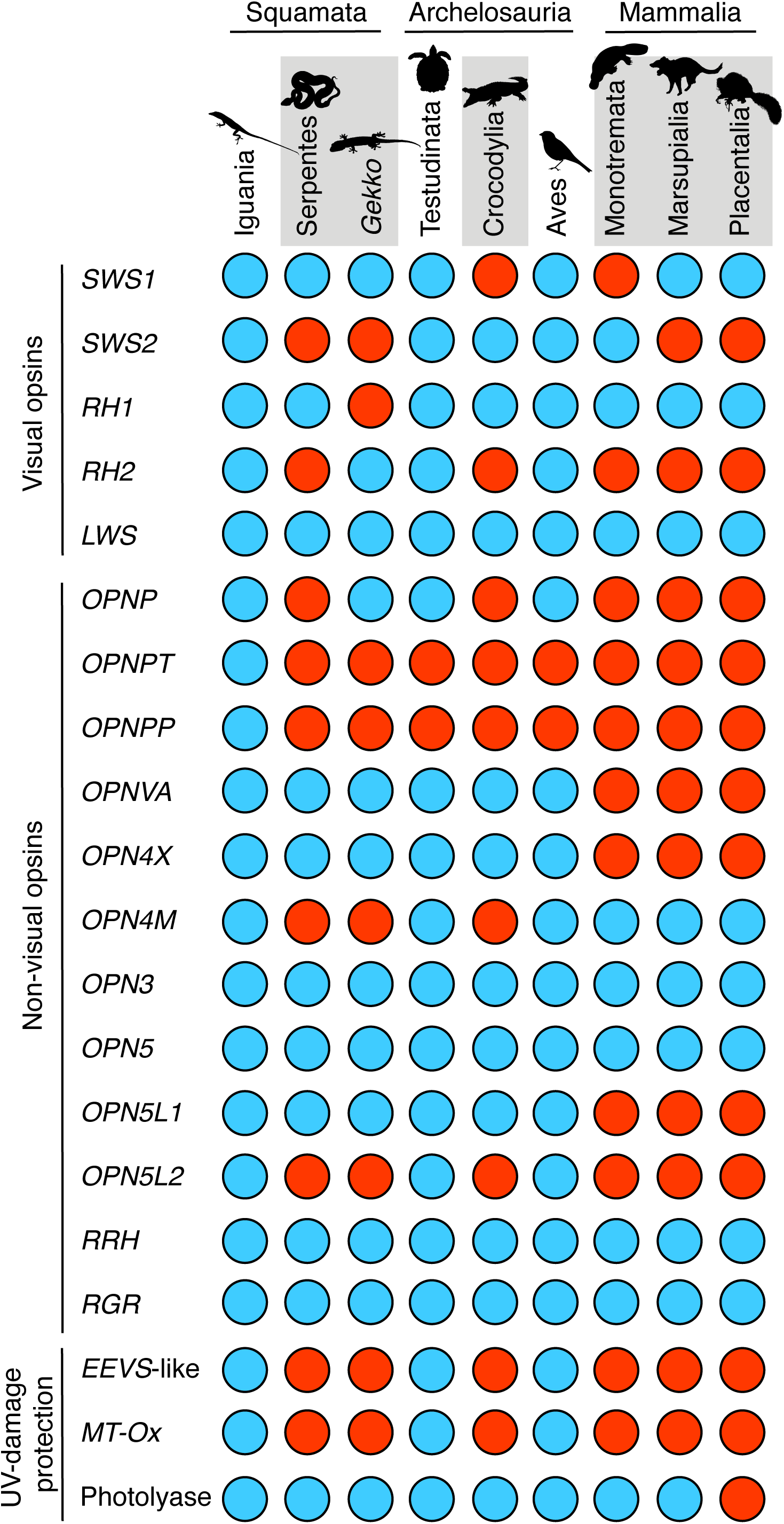
Temporal distribution of serpent and gekkotan gene losses. Bars indicate gene inactivation dating estimates based on an average of estimates from four model assumptions. Topology and divergence times of extant taxa based on Zheng and Wiens (2016) and positions of extinct taxa (†) from Martill et al. (2015). Taxa of focal species indicated in red. Bars indicate average point estimates for gene inactivation dates (ESM, Table S5). Colored branches indicate uncertainty with timing of gene loss due to whole gene deletions or ambiguous dN/dS ratios (*TAS1R;* see text and ESM, Table S4). Note that an additional three gene losses in *Gekko* (*OPNPT*, *OPN5L2*, *EEVS-like*), based on negative BLAST results and synteny data, are not shown in the figure. Silhouettes from phylopic.org (ESM, Table S7).

EGA’s annotations for predicted genes in squamates do not include the claw keratin *HB1*. Instead, the Anolis query sequence for *HB1* (EU755366, keratin aHB1 mRNA) has 100% query cover and identity to an *Anolis* mRNA described as “keratin, type II cytoskeletal cochleal” (NM_001293121). The GenBank entry indicates that this reference sequence is the claw keratin gene *HB1* described by Eckhart et al. (2008), suggesting that, despite the annotation, this mRNA encodes *HB1*. Whereas *Gekko japonicus* has a predicted sequence annotated as “keratin, type II cytoskeletal cochleal-like” with 100% query cover and 80% similarity, all of the serpent sequences with similar annotations have a much lower query cover (55-59%) and identity (69-70%). These values for the snake sequences are almost identical to those of an *Anolis* predicted sequence also annotated as “keratin, type II cytoskeletal cochleal” (60% query cover, 70% identity), implying that these predicted snake sequences represent a gene distinct from *HB1*. Consistent with this hypothesis, BLASTing *HB1* sequence probes against WGS yielded only negative BLAST results for snakes (ESM, Table S2), indicating gene loss in this clade. Synteny was difficult to establish given that *HB1* in *Anolis is* flanked by *ETAA1* (5’ end) and two keratins (3’ end) that EGA does not predict to be present either in snakes or *Gekko* (keratin, type II cuticular Hb4) or snakes alone (keratin, type II cuticular Hb5). Therefore, I tentatively conclude that *HB1* has been deleted from snake genomes, but further analyses will be required to confirm this.

### 3.2. Gustatory genes

In contrast to crocodylians, testudines, and non-serpent squamates, analyses of gustatory genes revealed the loss of numerous taste receptor types across the snakes surveyed (ESM, Tables S2 and S3). *TAS1R1* returned negative BLAST results from both EGA and WGS for all of the snakes surveyed. *TAS1R1* is flanked by *NOL9*and *ZBTB48*in sauropsids (ESM, Table S2), and both genes were found on the same contig in *Vípera berus*, *Protobothrops mucrosquamatus*, and *Ophiophagus hannah*, implying that *TASlRl* was deleted from their genomes. *Python bivittatushas* intact copies of TAR? and *TAS1R3*, whereas the colubroids in this study lack functional copies of both genes: *TAS1R2* is a pseudogene and *TAS1R3* returned negative BLAST results in all seven species. The genes that flank *TAS1R3* in amniotes (*CPTP*, *DVLT*) are located on the same contig in *Vípera berus* and *Ophiophagus hannah*, providing strong evidence of whole gene deletion in these taxa. All of the colubroids show evidence of deletion of exon 3 in *TAS1R2*, suggesting this gene may have been lost in their MRCA. However *TAS1R2* has numerous unambiguous inactivating mutations shared by viperids, but none shared by the elapid *Ophiophagus hannah* and/or the colubrids *Thamnophis sirtalis* and *Pantherophis guttatus*, raising the possibility that loss of *TAS1R2* occurred in parallel. Modeling the evolution of *TAS1R2* estimated that non-colubroid branches were under strong purifying selection (model 6 ω = 0.1677-0.2817), whereas crown colubroid branches ranged from strong to relaxed purifying selection (model 5 ω = 0.1182-0.8663; ESM, Table S4), rather than strictly relaxed selection. This is a bit difficult to interpret, given that crown Viperidae has an estimated dN/dS ratio ranging from 0.4112-0.666, despite the expectation of relaxed selection following the shared inactivating mutations. This implies that even after loss-of-function mutations accrued in this gene, there remains some selective constraint. For the bitter receptors, I only examined the three snake species in the EGA pipeline. *Python bivittatus*, *Protobothrops mucrosquamatus and Thamnophis sirtalis* are predicted to have two, one and zero bitter receptors, respectively (ESM, Table S3), which are the lowest copy numbers predicted across the reptiles I examined, including two alligators (4-7 TAS2Rs), three testudines (6-13 TAS2Rs), *Gekko* (6 TAS2Rs) and *Anolis* (28 TASRs). The only taste receptor gene not absent in any of the snakes is *PKD2L1* (ESM, Table S2), which encodes the sour taste receptor.

### 3.3. Light-associated genes

Three of the five visual opsin photopigment genes were intact in snakes: *SWS1* (short wavelength-sensitive opsin 1), *RH1* (rod opsin), and *LWS*(long wavelength-sensitive opsin). *SWS2* is a pseudogene in all eight focal species, with numerous loss-of-function mutations in each snake (ESM, Table S2). All eight species share four inactivating mutations, suggesting loss of *SWS2 in* their MRCA. Remnants of *RH2* (*exon* 2) were only discovered in *Python bivittatus*, with EGA and WGS returning negative BLAST results in the remaining seven snakes. Multiple inactivating mutations confirm that this gene is nonfunctional *in P. bivittatus* (*ESM*, Table S2). The genes that flank *RH2* in other sauropsids (*MLN*, *GRM4*) were found in the snake assemblies, however both genes never occurred on the same contig.

Four light-associated genes could not be recovered through BLAST searches (ESM, Table S2), including opsin genes (*OPNP*, *OPNPT*, *OPN5L2*) and a UV-light damage prevention gene (*EEVS*-like). The genes flanking *OPNP* in archelosaurians (*D0C2B*, *TEX14*) were not found to occur on the same contig of any of the snake assemblies, preventing confirmation of gene deletion. *PLEKHG7* overlaps with intron 1 of *OPNPT* in *Anolis*, and this gene was found in the snake assemblies, providing additional evidence that *OPNPT* was deleted. The genes flanking *OPN5L2* in sauropsids (*CDC5L*, *MUT*) are located on the same contig in *Ophiophagus hannah*, providing positive evidence of gene deletion. Finally, *MT-Ox* and *MITF* flank *EEVS*-like in vertebrates (Osborn et al., 2015), and both are on the same contig in *O. hannah*, again indicating whole gene deletion.

*OPN4M* returned negative BLAST results for all of the snakes except for *Python bivittatus*, which retains exons 5, 6 and 8 with multiple inactivating mutations (ESM, Table S2). While snakes retain the genes flanking *OPN4M* in tetrapods ( *WAPL*, *LDB3*), in no cases were these genes found on the same contig. Two additional genes, *OPNPP* and *MT-Ox*, were recovered as pseudogenes in all eight snakes. I found exon 1 of *OPNPP*, with 1-bp and 14- bp deletions shared by *P. bivittatus* and colubroids, suggestive of inactivation in their common ancestor. *MT-Ox*, another UV-protection enzyme gene, was similarly inactivated in all eight snakes, with 12 loss-of-function mutations shared by *P. bivittatus* and colubroids.

I was able to perform gene inactivation estimates for the following light-associated pseudogenes in serpents (ESM, Table S5): *RH2* (*mean\* 160.4 Ma; range: 129.9-184.6 Ma), *OPN4M* (*159* Ma mean; 141-173.7 Ma range), *MT-Ox* (143.3 Ma mean; 137.9-148.69 Ma range) and *SWS2* (*mean:* 110.6 Ma; range: 103.7-118.3 Ma). The dN/dS ratio estimates for *OPNPP* on the stem snake branch exceeds 1.0 using two different codon models (F1X4: 12.4771; F3X4: 999), though neither model is significantly differently from 1.0 (p > 0.25) suggesting the gene has been under relaxed selection on this branch.

As a positive control for patterns of light-associated gene loss, I examined the same set of genes in a member of the historically nocturnal gecko lineage (*Gekko japonicus*). Among visual opsins, Liu et al. (2015) previously reported that both *RH1* and *SWS2* are inactivated in *G. japonicus*, both of which I confirmed. I additionally found evidence of *LWS* inactivation, based on a single premature stop codon in exon 4 (ESM, Table S2, Dataset S1). dN/dS ratio branch test analyses of *LWS* indicates an elevated ratio for the *G. japonicus* branch (ω = 0.2086-0.2525) relative to the background (ω = 0.0902-0.1204), but the fit for this model was not significantly different from a one ratio model. If this gene is indeed nonfunctional, the single putative inactivating mutation indicates that this gene was pseudogenized after its split from *Gekko gecko* (Kojima et al., 1992). Beyond the visual opsins, I found evidence of gene loss in six additional light-associated genes (ESM, Table S2): *OPNPP* (inactivating mutations), *OPNPT* (negative BLAST results), *OPN4M* (inactivating mutations), *OPN5L2* (negative BLAST results), *EEVS-like* (negative BLAST results, confirmed via synteny), and *MT-Ox* (inactivating mutations). By contrast, all of these genes are intact in the diurnal *Anolis carolinensis* (ESM, Table S2). Gene inactivation dating analyses gave estimates for the loss of some genes on the stem Gekkotan branch (*SWS2:* 184.6 Ma [ω = 1.2716-1.9877]; *OPNPP*: 173 Ma mean, 202.1-136.6 Ma range; *OPN4M:* 152.3 Ma mean, 168.4-134.3 Ma range) and others within gekkonids (*MT-Ox*. 87.1 Ma mean, 105.1-69.6 Ma range; *RH1:* 80.9 Ma mean, 92.2-69.7 range; Fig. 1; ESM, Tables S4 and S5).

## 4. Discussion

### 4.1. Claw keratins

Analyses of hard keratins in *Anolis carolinensis* (Carolina anole; Iguania) have uncovered two putatively claw-specific keratins (Eckhart et al., 2008; Alibardi et al., 2011): HA1 (hard acid keratin 1) and HB1 (hard basic keratin 1). Further analyses by Dalla Valle et al. (2011) found that HA1 is also expressed in the claw-forming epidermis of the lacertoid *Podarcis sicula*, suggesting claw-specific HA1 expression may be widespread within Squamata. PCR amplification similarly found intact copies of the gene encoding HA1 in another iguanian (*Iguana iguana*), an anguimorph (*Varanus gouldii*), two gekkotans ( *Tarentola mauritanica*, *Hemidactylus turcicus*), and even the tuatara (*Sphenodon punctatus*), whereas it is a pseudogene in the legless anguimorph slow worm (*Anguis fragilis;* Dalla Valle et al., 2011), providing additional evidence of claw-specificity.

Although PCR amplification by Dalla Valle et al. (2011) failed to find *HA1* in the ball python (*Python regius*) and the corn snake (*Pantherophis guttatus*), I was able to obtain exon 1 of *HA1* in a pythonid, four viperids and an elapid. Multiple loss-of-function mutations, including a shared 8-bp insertion, indicate that this gene was lost in a common ancestor of the study species. Similarly *HA2* appears to be deleted from the genomes of these snakes. Given the absence of claws in snakes, these data provide further evidence that these keratins are claw-specific.

I further tested this hypothesis by estimating the timing of *HA1* inactivation and comparing it to evidence of limb loss in the serpent fossil record. Although all extant snakes are limbless, at least some snakes from the Late Cretaceous, including simoliophiids (Caldwell and Lee, 1997; Rage and Escuillié, 2000; Tchernov et al., 2000) and *Najash* (Apesteguía and Zaher, 2006), retained hind limbs, and *Tetrapodophis*, a putative snake from the Early Cretaceous, had four legs (Martill et al., 2015). While *Tetrapodophis* and *Najash* are typically reconstructed as stem serpents (Apesteguía and Zaher, 2006; Zaher et al., 2009; Martill et al., 2015; Pyron, 2016), phylogenetic reconstructions using phenotypic datasets (Apesteguía and Zaher, 2006; Zaher et al., 2009; Longrich et al., 2012; Hsiang et al., 2015; Pyron, 2016), combined molecular and phenotypic datasets (Hsiang et al., 2015; Reeder et al., 2015; Pyron, 2016) and morphological datasets with molecular constraints (Martill et al., 2015) have recovered simoliophiids, and even *Najash* (Reeder et al., 2015; Pyron, 2016), as nested within crown Serpentes, suggesting that limb loss occurred at least twice within snakes. Using a molecular dating method (Meredith et al., 2009) and assuming divergence dates from Zheng and Wiens’ (2016) phylogenomic timetree, I estimated that *HA1* was inactivated 113 Ma (point estimate; range: 108-118.5 Ma; Fig. 1; ESM, Tables S4 and S5), which pre-dates Zheng and Wiens’ (2016) estimate for crown Alethinophidia (92.7 Ma) but post-dates their age for crown Serpentes (128.1 Ma). *HA1* inactivation on the stem alethinophidian branch is consistently reproduced with eight alternative timetrees (Pyron and Burbrink, 2014; Hsiang et al., 2015; Pyron, 2016; ESM, Table S6) and four model assumptions (32 estimates), supporting the hypothesis that two or more snake lineages lost their limbs in parallel.

While an inactivation date of 113 Ma is coeval with the upper boundary of the stem serpent *Tetrapodophis* (Aptian; 125-113 Ma), it pre-dates the occurrence of the limbed simoliophiids by at least 13 million years (Cenomanian; 100.5-93.5 Ma). There are number of possible explanations for this apparent discrepancy. First, assuming that simoliophiids are not direct ancestors of any alethinophidians, there may be a ghost lineage extending to earlier than the Cenomanian, predating the gene inactivation estimate. Second, the reduced limbs of simoliophiids may not actually have had claws, such that claw loss preceded limb loss. Consistent with this hypothesis, simoliophiids had internal limbs with respect to the ribcage, and therefore were unlikely to retain claws, whereas the probable stem-serpent *Najash* had limbs that were external to the ribcage (Zaher et al., 2009). Finally, gene inactivation dating estimates are contingent upon the timetree used (ESM, Table S6), and timetree divergence estimates are largely influenced by the fossil calibrations implemented. As such, future analyses may benefit from exploring additional robust calibrations for snakes.

### 4.2. Gustatory genes

Recent research on the molecular basis of gustation in vertebrates has identified proteins associated with specific tastes (Chandrashekar et al., 2006; Yarmolinsky et al., 2009). TAS1R1+TAS1R3 form a heterodimeric taste receptor involved in umami taste, binding tastants including L-amino acids and nucleotides, whereas TAS1R2+TAS1R3 heterodimers bind sweet tastants, such as sugars. TAS2Rs form a family of bitter taste receptors, which detect a variety of potentially toxic compounds. Whereas sweet, umami and bitter taste proteins are G protein-coupled receptors, sour taste is conferred by a transient receptor potential ion channel known as PKD2L1. The identity of the salt receptor remains contentious, and therefore was not included in this study. All of these taste receptors are localized to taste buds within papillae of the tongue and/or the soft palate, and provide animals with a variety of information on nutritional content when ingesting food (Chandrashekar et al., 2006; Yarmolinsky et al., 2009).

Whereas taste buds appear to be widespread in squamates (Schwenk, 1985), snakes are reported to completely lack them (Schwenk, 1985; Young, 1997). However, the absence of taste buds within snakes is not without controversy (reviewed in Young, 1997), with contradictory results reported even within the same species (Kroll, 1973; Young, 1997; Berkhoudt et al., 2001). Because genomic data have the potential to validate the retention or loss of gustation in snakes, I searched for umami, sweet, bitter and sour taste receptor genes in the serpent genomes and compared the distribution of retained genes to several reptilian outgroups.

In contrast to crocodylians, testudines, and two non-serpent squamates, snakes show an extensive pattern of taste receptor loss. I found evidence that umami taste was lost at least since the MRCA of pythonids and colubroids, bitter taste is highly minimal to absent in *Python bivittatus*, *Protobothrops mucrosquamatus and Thamnophis sirtalis*, and at least some colubroid lineages have lost sweet taste, possibly in parallel. By contrast, the sour taste receptors appear to be intact in all of the species queried. While taste appears to have been de-emphasized during serpent history, here I provide genomic evidence that the capacity for at least some tastes are retained in viperids, elapids, colubrids and especially pythonids, consistent with some reports of taste buds in snakes (e.g., Kroll, 1973; Berkhoudt et al., 2001). Nonetheless, there are clearly inconsistencies in the data (Young, 1997) suggesting that when present, they may be very few in number, and as such taste may be a relatively unimportant sensory modality in this taxon.

### 4.3. Light-associated genes

Based on comparative anatomy, the eyes of early snakes lost a number of traits present in the squamate ancestor, including the annular pad of the lens, sclerotic rings, scleral cartilage, iris muscles and cone oil droplets (Walls, 1942; Atkins and Franz-Odendall, 2016). This ocular degeneration has historically been interpreted as the result of stem snakes shifting to a scotopic niche, similar to that inferred from the ocular anatomy of mammals and crocodylians (Walls, 1942). Genes encoding visual pigments and associated loci have also been lost/inactivated in a number of vertebrates adapted to scotopic niches, including nocturnal, fossorial and deep-sea marine species (Yokoyama et al., 1999; Kim et al., 2011; Jacobs, 2013; Meredith et al., 2013; Gerkema et al. 2013; Emerling and Springer, 2014; Fang et al., 2014; Emerling and Springer, 2015; Springer et al., 2016; Emerling, 2016)

Concomitant with the regression of some features of serpent eyes, I found that a number of genes encoding proteins with light-associated functions were inactivated or lost in snakes (ESM, Table S2). Consistent with previous microspectrophotometric, genomic, and retinal mRNA studies (Sillman et al., 1997; Sillman et al., 1999; Sillman et al., 2001; Davies et al., 2009; Hart et al., 2012; Castoe et al., 2014; Simoes et al., 2015; Simoes et al., 2016a, b), three of the five visual opsin genes were intact in snakes (*SWS1RH1*, *L WS*), whereas *SWS2* is pseudogenized in all eight species and remnants of *RH2* were only recovered in *Python bivittatus*. The presence of S2 and *RH2* pseudogenes confirms that these opsins are not simply being expressed at low levels in the retina or co-opted for non-visual functions. Although the absence of ÆHpseudogenes in the colubroids prevents confirmation of loss in their MRCA, both the mean (160.4 Ma) and range (129.9-184.6 Ma) of gene inactivation estimates for *RH2* (*ESM*, Table S5) predate Zheng and Wiens’ (2016) divergence estimate for crown Serpentes (128.1 Ma), suggesting loss of *RH2* occurred in a stem snake (Fig. 1). By contrast, *SWS2* inactivation point estimates (mean: 110.6 Ma; range: 103.7-118.3 Ma; ESM, Table S5) postdate the crown Serpentes divergence and predate the origin of crown Alethinophidia (92.7, Ma), signifying parallel losses of WS in alethinophidians and scolecophidians (Simoes et al., 2015). If confirmed with additional genomic sequences, this implies that the common ancestor of snakes had trichromatic color vision, similar to humans, as opposed to the dichromatic color vision previously implied by parsimony.

A number of light-associated genes with non-image forming functions were deleted/pseudogenized. Among these genes were some encoding light-sensitive pigments expressed in the pineal gland (OPNP, pinopsin; Okano et al., 1997; Frigato et al., 2006; Su et al., 2006; Wada et al., 2012), the squamate parietal (“third”) eyes (OPNPT, parietopsin; Su et al., 2006; Wada et al., 2012; OPNPP, parapinopsin; Wada et al., 2012), adrenal glands (OPN5L2; Ohuchi et al., 2012), and a variety of tissues (OPN4M, mammal-like melanopsin; Bellingham et al., 2006) of vertebrates. Additionally, genes encoding two enzymes involved in synthesizing the endogenous UV-protectant sunscreen gadusol were deleted/pseudogenized in the snakes examined (*EEVS-like*, *MT-Ox*; Osborn et al., 2015). Shared inactivating mutations in *OPNPP* and *MT-Ox*, plus estimates of gene inactivation timing for *OPN4M* (159 Ma) and *MT-Ox* (*143.3* Ma) that predate Zheng and Wien’s (2016) date for crown Serpentes (128.1 Ma), point to additional light-associated gene losses in a stem serpent.

Castoe et al. (2014) was unable to find copies of the visual and non-visual opsin genes in *Python bivittatus*, and Osborn et al. (2015) reported the absence of EE-like and Mr-x orthologs in both P. *bivittatus*and *Opiophagus hannah*. Using synteny data and the identification of pseudogenic remnants of these genes (ESM, Table S2, Dataset S1), I confirmed that the absence of most of these genes is not attributable to assembly errors, and extends from pythonids to viperids, elapids and colubrids, indicating widespread loss of light-associated genes in snakes.

This pattern of light-associated gene loss is highly similar to that seen in mammals (Gerkema et al., 2013) and crocodylians (Emerling, 2016; Fig. 2), two lineages thought to have experienced ‘nocturnal bottlenecks’. Geckos are similarly thought to have undergone a long period of scotopic adaptation, based on both ocular modifications (Walls, 1942) and the phylogenetic distribution of activity patterns in this clade (Gamble et al., 2015). In addition to the pseudogenization of *SWS2* and *RH1* (Liu et al., 2015), I found evidence of gene loss in four non-visual photopigment genes (*OPNPP*, *OPNPT OPN5L2*, *OPN4M*) and the two gadusol synthesis genes (*EEVS*-like, *MT-Ox;* ESM, Table S2). The convergent patterns of gene pseudogenization/loss, in genes with clear functional associations to light, both in snakes and three historically nocturnal lineages corroborate evidence from ocular anatomy that snakes were dim light-adapted early in their history.

**Figure 2.**
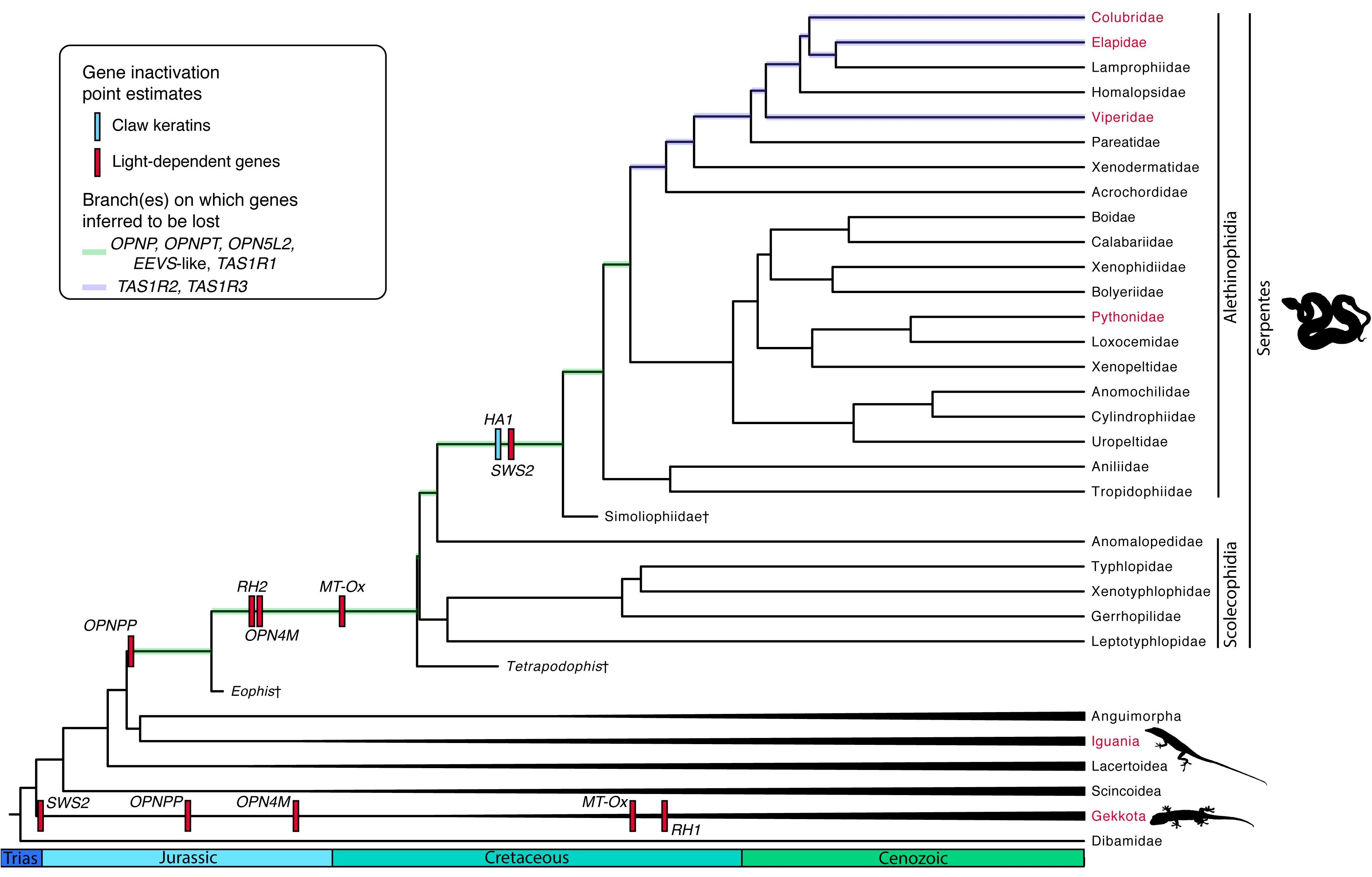
Comparisons of light-associated gene losses across amniotes. Blue circles = gene intact in common ancestor of clade; red circles = gene inactivated/deleted in common ancestor of clade. Clades associated with scotopic adaptations early in their respective histories indicated by shading. Silhouettes from phylopic.org (ESM, Table S7).

### 4.4. Genomic regression and the ancestral ecology of snakes

The question of the ecological conditions under which snakes first evolved is a contentious issue for serpent evolutionary biologists. Some researchers have favored a marine origins hypothesis, which postulates that a terrestrial stem-snake became secondarily-aquatic. Evidence for this hypothesis comes from phylogenetic reconstructions that suggest that the extinct marine mosasaurs are sister to snakes (Lee et al., 1999; Lee, 2005; Reeder et al., 2015; though see Pyron [2016] for conflicting placements), the discovery of Cretaceous marine snakes (Simoliophiidae) that retain diminutive hindlimbs (Caldwell and Lee, 1997; Rage and Escuillié, 2000; Tchernov et al., 2000), and phenetic and parsimony analyses that found serpent eyes to be similar to those of fishes (Caprette et al., 2004). Although an explanation for the adaptive basis of limb loss is rarely offered, presumably it would have reduced friction for early swimming serpents. One piece of evidence for the marine origins hypothesis that appears to have gone unnoticed is that snakes have had a massive reduction in their taste buds (Young, 1997), a condition also found in secondarily aquatic cetacean mammals (reviewed in Feng et al., 2014). The condition in cetaceans has been hypothesized to be due to the masking of tastants diluted in overwhelmingly saline marine environments (Feng et al., 2014), which may also explain the condition in snakes.

A second common hypothesis for the origin of snakes posits that the earliest species were fossorial. Evidence for this comes in part from the phylogenetic distribution of burrowing snakes, many of which are among the most basal serpent lineages (e.g., Scolecophidia, Aniliidae, Tropidophiidae, Cylindrophiidae, Anomochilidae, Uropeltidae; Zheng and Wiens, 2016), possibly indicating retention of an ancestral state in these taxa. Many squamates with reduced or absent limbs are burrowers, including dibamids, amphisbaenians, and some scincids and gymnophthalmids (Wiens et al., 2006), as are most caecilian amphibians, implying that limb loss is a common consequence of fossoriality. Using principal components analyses, Wiens et al. (2006) found that limbless squamates generally fall into two ecomorphs: long-tailed surface dwellers and short-tailed burrowers. Notably, snakes typically group with short-tailed burrowers, regardless of ecology. Among fossils, the inner ear vestibule in the stem serpent *Dinilysia* indicates that it had a morphology most similar to burrowing squamates (Yi and Norell, 2015), and other traits, such as the low neural arches in the stem snakes *Najash* (Apesteguia and Zaher, 2006) and *Coniophis* (Longrich et al., 2012), have been interpreted as indicative of a fossorial lifestyle.

Comparisons of ocular anatomy indicate that the visual system of snakes has undergone a great deal of regression (Walls, 1942), likely in association with adaptation to a scotopic niche. Inhabiting dimly-lit marine ecosystems and dedicated burrowing may lead to similar phenotypes, and unsurprisingly both have been invoked as possible selective pressures to snakes’ regressed visual system (e.g., Walls, 1942; Caprette et al., 2004). However, an extended commitment to nocturnality can also lead to the loss of various vision-related traits, and such a hypothesis has been invoked for both ancestral mammals and crocodylians (Walls, 1942; Gerkema et al., 2013; Emerling, 2016) to explain the loss of numerous ocular structures. Accordingly, extensive nocturnality may be sufficient to lead to the regressed eye anatomy in extant snakes. In fact, a great number of snakes, including many of the more basal lineages, are nocturnal, and Hsiang et al. (2015) reconstructed serpent evolutionary history as being dominated by a nocturnal activity pattern. Regression of genomic loci can weigh in on the question of snake origins by comparing patterns of gene loss to other taxa and estimating the timing of gene inactivations.

Taste receptor loss has occurred frequently in the evolutionary history of amniotes, and is typically assumed to be associated with changes in diet (Zhao et al., 2010; Zhao et al., 2012; Jiang et al. 2012; Hayakawa et al. 2014; Liu et al. 2016), though predictions of gene loss based on diet are not always straightforward (Feng and Zhao, 2013). The degree of taste receptor loss that I report here in colubroids has so far only been documented in cetaceans (Feng et al., 2014), which have also lost the umami, sweet and bitter taste receptors. Whales have additionally inactivated the sour receptor, suggesting an even greater reduction in gustation than colubroids. Feng et al. (2014) discussed several hypotheses for taste receptor gene loss in cetaceans, and included the possibility that tastants would be difficult to detect while diluted in highly saline water. Marine pinnipeds have also lost their sweet and umami taste receptors (Jiang et al., 2012) and have had extensive losses of bitter taste receptors (Liu et al., 2016), providing additional support for this hypothesis. The pervasive loss of taste in marine mammals and colubroids may be suggestive of marine adaptations in Serpentes, but would seem to indicate that more crownward taxa, rather than stem snakes, might have been marine adapted. However, I found that the marine turtle *Chelonia mydas* retains umami, sweet, sour and six bitter receptors (ESM, Tables S2 and S3), implying that inhabiting a marine habitat may be insufficient to lose gustation channels. An alternative hypothesis is that the reduction of taste buds and taste receptors in snakes is not related to diet, but instead is associated with a greater forking of the tongue for vomeronasal sensation (Schwenk, 1994). As additional evidence for this hypothesis, the only other squamates with tongues that are as strongly forked as snakes are varanids, which also appear to completely lack taste buds (Schwenk, 1985; Young, 1997). Perhaps as the tongues of these clades became more strongly associated with signaling gradients of molecules to the vomeronasal system, there was a trade-off that led to a reduction in gustatory function.

The loss of light-associated genes in snakes would seem to have some bearing on the question of serpent origins. The pattern of gene loss is highly convergent with *Gekko japonicus*, crocodylians and mammals (Fig. 2), three lineages thought to have independently undergone a lengthy period of nocturnal adaptation (Walls, 1942; Gerkema et al., 2013; Gamble et al., 2015; Emerling, 2016): (1) snakes, *Gekko*, crocodylians, monotremes and therian mammals have each lost two of the five classes of vertebrate-specific visual opsins; (2) snakes, crocodylians and mammals lost the pineal-expressed pinopsin; (3) snakes, *Gekko*, crocodylians and mammals lost two opsins (parietopsin, parapinopsin) putatively associated with the parietal eye; (4) snakes, *Gekko and* crocodylians lost mammal-like melanopsin (OPN4M), whereas mammals lost the paralogous *Xenopus-like* melanopsin (OPN4X); (5) snakes, *Gekko*, crocodylians and mammals lost OPN5L2; and (6) snakes, *Gekko*, crocodylians and mammals lost two enzymes responsible for synthesizing the sunscreen compound gadusol (EEVS-like, MT- Ox). Though the loss of parietopsin and parapinopsin may be linked to scotopic adaptation, previous research (Borges et al., 2015; Emerling, 2016) has shown that they were also lost in turtles and birds, two lineages that lack a parietal eye and are not traditionally thought to have experienced an extensive period of scotopic adaptation. Instead, the loss of *OPNPP* and *OPNPT* may be more broadly associated with adaptations to thermal stability (see discussion in Emerling, 2016). The remaining light-associated genes are broadly retained in vertebrates (Tomonari et al., 2008; Gerkema et al., 2013; Osborn et al., 2015; Emerling, 2016). Notably, they are typically found intact in the genomes of fishes and turtles, including the marine-adapted *Chelonia mydas*, indicating that inhabiting a marine habitat is unlikely to have led to the loss of these genes. Importantly, this neither implicates nor rules out fossoriality as the basis for light-associated gene loss in snakes, but given that fossoriality can lead to the extensive loss of genes associated with vision (Kim et al., 2011; Emerling and Springer, 2014; Feng et al., 2014), this remains a plausible hypothesis. Future studies would benefit from investigating the patterns of non-visual opsin and UV-protection gene loss in fossorial species and contrasting it with nocturnal taxa.

The mean inactivation date estimates for *RH2* (160.4 Ma), *OPN4M* (159 Ma), and *MT-Ox* (143.3 Ma) in snakes predate the loss of limbs and the estimated date of *HA1* inactivation by 30+ million years. This may point to an extended period of fossoriality before complete limb loss occurred, or that early snakes were highly nocturnal and fossoriality or marine adaptations came later in their evolutionary history. It is important to note that these gene inactivation estimates assume that pseudogenization along a branch involves a transition from purifying selection to relaxed selection (Meredith et al., 2009). Though a very brief period of positive selection favoring the loss of a gene likely would not greatly distort these estimates, an extensive period of positive selection could. A simpler test of the hypothesis that scotopic adaptations presaged limb loss would involve comparisons of alethinophidian and scolecophidian genomes. If scolecophidian and alethinophidian genomes share inactivating mutations in light-associated genes, but not *HA1*, this would further indicate that scotopic adaptation preceded limb loss in snakes. Recent discoveries that snakes have mutations, including deletions, of numerous limb-specific enhancers (Infante et al., 2015; Leal and Cohn, 2016) provide additional loci for comparison to determine the timing of limb loss in snakes.

### 4.5. The importance of light-dependent regressed traits in understanding serpent history

As discussed above, the convergent patterns of light-associated gene loss seem to implicate nocturnality as a more plausible explanation than marine adaptation, given the retention of these genes in the green sea turtle. In addition, some morphological traits involved in light perception show patterns of regression consistent with nocturnality and/or fossoriality but not with marine adaptation. Snakes lack the carotenoid-based cone oil droplets typical of diurnal sauropsids (Walls, 1942). These droplets filter wavelengths in order to sharpen the spectral peaks of cone opsin pigments, thereby minimizing spectral overlap and increasing hue discrimination. However, this filtering function also leads to diminished sensitivity (Vorobyev, 2003), thereby rendering them less adaptive under scotopic conditions. Among squamates, cone carotenoids have also been lost in nocturnal (geckos [Gekkota]; beaded lizards [Helodermatidae]) and secretive (night lizards [Xantusiidae]) species (Walls, 1942), with the droplets being completely lost in some gecko lineages (Röll, 2000). Both placental mammals and crocodylians have disposed of oil droplets completely, with monotremes and marsupials lacking the carotenoid pigments, consistent with the ‘nocturnal bottlenecks’ hypothesized for these taxa. Walls (1942, pg. 203) reported that “fossorial lizards such as *Anniella* have lost most of the pigment of the oil-droplets”, though does not elaborate on other species. By contrast, among marine-adapted sauropsids, sea turtles (Granda and Haden, 1970) and penguins (Bowmaker and Martin, 1985) retain carotenoid oil droplets.

Snakes have also lost sclerotic rings (Walls, 1942), bony elements in the orbit that are associated with the accommodation of the eye (Atkins and Franz-Odendaal, 2016). Accommodation is more typical of photopic vision, during which the high acuity cone photoreceptors are functioning and allow for focusing on objects of different distances. Sclerotic rings have been lost in mammals and crocodylians, further evidence of the influence of extended nocturnality on the visual system. An extensive survey of squamates (Atkins and Franz-Odendaal, 2016) demonstrated that sclerotic rings were additionally lost in two legless, fossorial clades (Dibamidae; Rhineuridae), indicating fossoriality may explain the loss of these elements as well. By contrast, sea turtles (Brudenall et al., 2008), marine iguanas (de Queiroz, 1982) and penguins (Suburo and Scolaro, 1990) retain sclerotic rings, as did numerous lineages of extinct marine reptiles, including mosasaurs (Yamashita et al., 2015), sauropterygians (Storrs and Taylor, 1996; Druckenmiller, 2002; Klein, 2009), ichthyosaurs (Motani et al., 1999), thalattosuchians (Nesbitt et al., 2013), and thalattosaurs (Müller, 2005). It is difficult to determine whether any marine sauropsids lost their sclerotic rings as an adaptation for aquatic vision, since the fragile ossicular constituents of the ring are not always prone to preservation. However, it is noteworthy that sea turtles maintain sclerotic rings despite a marine history that may date back to the Early Cretaceous (Cadena and Parham, 2015).

These light-associated traits do not alone rule out the marine origins hypothesis in favor of a nocturnal and/or fossorial scenario for early snakes, but they do highlight the importance of comparative analyses in inferring evolutionary history. Additional morphometric analyses, similar to those of Wiens et al. (2006) and Yi and Norell (2015), and comparisons of regressed anatomical and genetic characters should be instrumental in further understanding snake evolution in future studies.

### 4.6. Conclusions

The expectation that genomic loci co-regress with anatomical traits allows for the testing of a diversity of hypotheses. I tested three such hypotheses associated with the origin of snakes: (1) the keratins HA1 and HA2 have claw-specific expression; (2) snakes lack the capacity for taste; and (3) the earliest snakes were adapted to dim-light conditions. I found evidence in support of hypotheses (1) and (3), with pseudogenic and deleted copies of the putative claw keratin genes and multiple genes encoding proteins with light-associated functions, but I rejected the hypothesis that snakes have completely lost their capacity for taste, given the retention of numerous taste receptors in some snakes. The patterns of gene loss also inform the ecological origins of snakes, with some evidence for dim light adaptations in stem snakes prior to leg loss, and most taste losses occurring in parallel within crown lineages. Both genomic and anatomical regression of traits, including limbs, taste and light-associated characters, may bring further clarity to the enigmatic origins of this diverse and ecologically disparate clade.

## Acknowledgements

I thank the Nachman Lab for helpful discussions. This research was supported by an NSF Postdoctoral Research Fellowship in Biology (Award #1523943). The *Thamnophis sirtalis* genome was produced by Richard K. Wilson and the McDonnell Genome Institute, Washington University School of Medicine

